# Identification of Thermotolerant Rice Genotypes with Allele Coding at Seedling Stage

**DOI:** 10.1101/2021.10.02.462852

**Authors:** Bandi Arpitha Shankar, Prashant Kaushik

## Abstract

Rice-The most important plant in the world to ensure food security. Heat is one of the main factors that greatly limit rice production. With the increasing global warming, industrialization there is a great effect on climate change which requires us to see various alternatives for strains that are more tolerant to heat so that some techniques are developed to filter a large number of genotypes for high temperature tolerance. Here we report the standardization of Temperature Induction Response (TIR) technique to identify thermotolerant rice genotypes. The phenotypic characteristics of Rice due to high temperature is calculated with germination (%), growth of the seedling and molecular analysis is also considered. The heat stress is provided to the plants with the help of TIR protocol with the adjustment of temperature to lethal (55°C) and sub-lethal levels (38-55°C) in a TIR chamber with alterations in humidity. Of the 74 genotypes screened, 14 showed thermo tolerance caused by high temperatures. Both tolerant and sensitive genotypes were separated based on their survival percentages. The tolerant class are selected based on the growth and development of genotypes having high survival percentage and also their shoot and root lengths, fresh and dry weights are compared to the heat tolerant checks N22, Dular and Nipponbare. These genotypes have intrinsic heat tolerance and thus can be explored as a source of donors in breeding programs intended for global warming. The molecular markers which are identified to be linked with heat tolerant class through allele code are quite helpful and can be used in marker assisted breeding approach to attain heat tolerance in cultivated varieties.

## INTRODUCTION

Rice is the most significant and essential cereal food grain domesticated all through the world particularly in Asia and Africa (Krishnan *et al*., 2011). The name wild rice is generally utilized for types of the genera Zizania and Porteresia, both wild and tamed, albeit the term may likewise be utilized for crude or crude assortments of Oryza (Lafarge *et al*., 2017). Rice, a monocot, is regularly developed as a yearly plant, albeit in tropical regions it can get by as a perennial and can deliver a ratoon crop for up to 30 years (Huang *et al*., 2012). Rice development is appropriate to nations and areas with low work expenses and high precipitation, as it is work concentrated to develop and requires sufficient water. The rice plant can develop to 1–1.8 m (3 ft 3 in–5 ft 11 in) tall, every so often relying upon the assortment and soil ripeness. It has long, slim leaves 50–100 cm (20–40 in) long and 2–2.5 cm (3/4–1 in) wide (Shi *et al*., 201; Kesh *et al*., 2021).

Rice enhancement for wetland rice fields is acknowledged to be answerable for 11% of the anthropogenic methane outpourings. Methane conveyed is achieved by long stretch flooding of rice fields cuts the earth off from ecological oxygen and causes anaerobic maturing of regular matter in the soil (Kesh and Kaushik, 2020). Methane creation from rice improvement contributes ~1.5% of complete anthropogenic nursery gases. Methane is on numerous occasions more impressive an ozone-exhausting substance than carbon dioxide (Yu *et al*., 2014).

A new report found that, due to rising temperatures and lessening sun-based radiation during the later extended lengths of the 20th century, the rice yield improvement rate has decreased in various bits of Asia, standing out from what may have been seen had the temperature and sun arranged radiation designs not happened (Kumar and Kaushik, 2021; Malhi *et al*., 2021). The yield improvement rate had fallen 10–20% at specific regions (Raza *et al*., 2019; Jain *et al*., 2018). The examination relied upon records from 227 farms in Thailand, Vietnam, Nepal, India, China, Bangladesh, and Pakistan (Jain *et al*., 2019). The instrument of this falling yield was not palatable, anyway may incorporate extended breath during warm nights, which devours energy without having the choice to photosynthesize (Fu *et al*., 2019).

In view of expanding temperatures by an unnatural weather change, plants are defenceless to intermittent warmth and dry spell pressure that generally influences the growth and development. Plants adjust to high temperature pressures with basal level resilience innate and can acquire resistance to serious temperature stress (Kim *et al*., 2011). Thermo tolerance acquired is very quick and has been demonstrated to be initiated during the phone’s acclimation until the temperature time frame is very high. The temperature influences the expansive range of cell and digestion parts, and outrageous temperatures force the seriousness of factors relying upon the degree of progress in temperature, power, and length (Kumar *et al*., 2021). The capacity to hold and change in accordance with the supra-ideal temperature results from both warmth harm avoidance and warmth touchy fix segments (Yufang *et al*., 2021). Seeds presented to the temperature of the sub-deadly prior to testing with weighty temperatures have a preferred development recuperation over the seeds tested straightforwardly to extreme temperatures (Satishraj *et al*., 2016). Both outrageous conditions (dry seasons and floods), on the off chance that they surpass certain basic periods, will have significant ramifications for rice and can cause complete disappointment of rice plants during delicate stages either as water deficiencies or exorbitant splashing (Chaturvedi *et al*., 2017). So, there is a need to receive a diverse methodology while considering the effect of high temperature stress, likewise zeroing in on other ecological pressing factors, which might be similarly adverse to rice usefulness (Priyanka et al., 2021b, 2021a). Resistance acquired for certain abiotic stress has been displayed to give cross security to different pressing factors like saltiness, cold temperatures, and dryness. Along these lines, assessing the overall exhibition of rice genotypes for high temperature resistance utilizing TIR approach (Liu *et al*., 2013).

The best temperature for rice germination is some place in the scope of 28 and 30 °C. High temperature impacts essentially all advancement periods of rice from germination to developing (Shah *et al*., 2011). The cut-off temperature at the seedling stage has been perceived as 35 °C; the essential sign of warmth stress is the vulnerable turn of events. By thinking about its significance, accessibility, uses and need for creation, heat pressure has acquired a substantial significance because of winning expanded temperatures (Burke *et al*., 2011). Warmth stress majorly affects every one of the phases of rice. By taking every single detail into thought, our investigation is focused on heat pressure lenient genotypes in rice accomplished by TIR convention so these genotypes can be additionally used to deliver heat open minded assortments either by reproducing program or by quality articulation examination (Sato *et al*., 2016).

## MATERIAL AND METHODS

### Experiment design

The area selected for the experiment is Phenotypic lab at Institute of Frontier Technology, Regional Agricultural Research Station, Tirupati, using TIR (Temperature Induction Response) protocol. The experimental material includes 74 diverse rice genotypes taken from Nellore, Maruteru, several land races and African lines (NERICA) including proven varieties for heat tolerance such as N22, Dular and Nipponbare which were used as genotype checks to choose tolerant sets (Prashanth *et al*., 2012). These TIR approaches involve first identification of challenging temperature and induction temperatures and then they are standardized before using the germplasms for intrinsic tolerance. Phenotyping of rice genotypes for thermo tolerance utilizing TIR protocol was set up in this lab and the same technique was utilized in this investigation.

Several genotypes were selected to carry out the work. A total of 74 genotypes which include wild relatives, land races, Aus groups, japonica lines etc., were selected for the work. The genotypes are tested for high temperatures to select heat tolerant lines along with proven check varieties.

### Treatments

Rice seeds were washed with distilled water 2-3 times and are stored for germination at room temperature. After 42 hours, the seedlings that attain 0.5 cm uniformly were selected and planted in an aluminium tray containing blotter paper moistened with water (Bado *et al*., 2016). Plates with these seeds are liable to sub-deadly (sub-lethal) temperatures (expansion in slow temperatures for each half an hour from 38°C to 55°C for 4 hours in this atmosphere - ‘Labline’ - (Humidity controlled chamber). Then these seeds are exposed to a deadly (lethal) temperature (55°C) (induced) for 2 hours. Sub-set of other seeds are exposed directly to the deadly temperature (non-induced). The seeds of the rice induced and non-induced are permitted to recover at room temperature for one week. A control tray is maintained at room temperature, which are not exposed to the temperature of the sub-lethal and deadly (Liu *et al*., 2017) conditions.

The treatment for the just sprouted seedlings is carried out in a special chamber called TIR (Thermal Induction Response) chamber where the plates are arranged according to the treatment. The chamber contains temperature adjustment along with humidity maintenance. The treatment varieties are tested along with the check varieties to compare the heat tolerance capacity in different varieties.

### Phenotypic Analysis

Highly vigour seedlings are selected as tolerant types because certain seedlings do not respond properly towards heat stress. The susceptible varieties did not germinate even under suitable conditions. Along with the germination the shoot and root lengths of seedlings are taken into consideration to select tolerant varieties. The maximum root length and shoot lengths of the growing seedlings are compared with each other and the highly tolerant, medium tolerant and susceptible varieties are segregated.

### Molecular Analysis

Out of 74, a set of 14 genotypes each under tolerant and control conditions were selected based on their survival percentage under sub-lethal conditions and are allowed for Selective line genotyping by comparing with the three checks i.e., N22, Dular and Nipponbare.

### DNA isolation and quantification

The tender, sprouted seedlings were maintained for 15 days at room temperature thereafter, the complete genomic DNA content of the seedlings was extracted using the CTAB method. The purified DNA pellet which was obtained through CTAB method is air–dried and dissolved in 50-100μl of TE (Tris base-1M, EDTA-0.5M) buffer (Tenorio *et al*., 2017). A proper DNA quantification was done using NanoDrop™ 1000 spectrophotometer (ThermoScientific, Wilmington, USA). Quantification of pure nucleic acid for DNA yielded a value of 260/280 ratio of ~1.80 which is proper for a DNA whereas the value more than 1.8 indicates the presence of RNA and less than 1.8 indicates the occupancy of proteins (Jagadish *et al*., 2012).

### Selection of primers

In General, 51 primers were chosen out of which 43 were SSRs reported for heat resilience and 8 were genic markers out of which 4 were brought to light from rice database www.gramene.org, and rest were designed from Primer 3 software. The microsatellite region of candidate genes were identified using SSRIT tool (http://archive.gramene.org/db/markers/ssrtool) and then primer designing was done using primer-3 v.4.0.0, a primer designing tool (http://bioinfo.ut.ee/primer3-0.4.0/) (Shanmughavadivel *et al*., 2017). Reported heat tolerant SSRs and genic SSRs used in the present study are provided in Table S1 and Table S2.

### PCR amplification and product isolation

The PCR amplification was performed using an Eppendorf Master cycler to understand the polymorphism. The reactions were performed with standard temperatures and are repeated for 35 times. Agarose gel was prepared to understand the detailed amplification with a permanent marker of 50bp or 100bp which was loaded along with the samples (Xie *et al*., 2014). For gel perception UV trans-illuminator was utilized and Alpha Innotech Multi Imager gel documentation framework program from Alpha Innotech, California, USA was used for photography (Vallejos *et al*., 2007).

## RESULTS

It was noted that all the 74 genotypes which were involved in sub-lethal temperatures were considered a major importance involving both tolerant and sensitive classes (Table 2). For all these genotypes the survival percentage was seen as a primary target, where all the 74 genotypes were segregated depending upon their survival ability and growth (Table 2). The survival percentages of all the genotypes were checked by taking their germination and growth into consideration. The survival percentage was calculated in two major classes i.e., SP between 80-100% and another one 60 % or less than 60% (Table 2). The genotypes that fall between the SP range of 80-100% were chosen as heat tolerant class and genotypes other than tolerant class were considered as heat sensitive class, depending upon their germination and growth after heat treatment in TIR chamber (Table 2).

**Table 1.** Genotypes selected for heat tolerance study.

| S.No. | Genotype | S.No. | Genotype | S.No. | Genotype |
| --- | --- | --- | --- | --- | --- |
| 1 | AC38460 | 27 | LN386 | 53 | NLR4002 |
| 2 | AC41038 | 28 | LN409 | 54 | NLR40024 |
| 3 | ARC10533 | 29 | Mahisugandh | 55 | Pokkali |
| 4 | Basmathi 370 | 30 | Minghui | 56 | Pusa1121 |
| 5 | Basmathi 386 | 31 | MTU1001 | 57 | Rajeshwari |
| 6 | Binuhungiri | 32 | MTU1010 | 58 | Ranbir Basmati |
| 7 | BPT1235 | 33 | MTU1061 | 59 | RNR150418 |
| 8 | BPT5204 | 34 | MTU1071 | 60 | Satya |
| 9 | Dalasaitha | 35 | MTU1121 | 61 | Shabigdhan |
| 10 | Dikhow | 36 | MTU3626 | 62 | Siddi 95 |
| 11 | Disang | 37 | Nagina22 (N22)* | 63 | Sona |
| 12 | Dular* | 38 | NBR16 | 64 | Sonasali |
| 13 | Erramallelu | 39 | Nilagiri | 65 | Swarna |
| 14 | FR 13 A | 40 | Nipponbare* | 66 | Swarna Sub1A |
| 15 | IR 1552 | 41 | NL1(*Og/Os*) | 67 | TKM-6 |
| 16 | IR 64 | 42 | NL61(*Og/Os*) | 68 | Udayagiri |
| 17 | Jagannadh | 43 | NL24(*Og/Os*) | 69 | Vajram |
| 18 | JGL 3844 | 44 | NL42(*Og/Os*) | 70 | Vasundhara |
| 19 | JGL 3855 | 45 | NLR145 | 71 | VL Dhan16 |
| 20 | Kab Aus R270 | 46 | NLR3042 | 72 | WGL347 |
| 21 | Kandagiri | 47 | NLR30491 | 73 | WGL482 |
| 22 | Kapilee | 48 | NLR3238 | 74 | WGL915 |
| 23 | Kolong | 49 | NLR3242 |  |  |
| 24 | Konark | 50 | NLR3354 |  |  |
| 25 | Koshihikari | 51 | NLR33671 |  |  |
| 26 | Lachit | 52 | NLR34242 |  |  |

\*, Check Varieties (reported as heat tolerant genotypes by earlier researchers)

**TABLE 2.** Survival percentage of genotypes under sub lethal conditions.

| S.No. | Genotype | SP/<br>SL*<br>(%) | S.<br>No. | Genotype | SP/<br>SL*<br>(%) | S.No. | Genotype | SP/<br>SL*<br>(%) |
| --- | --- | --- | --- | --- | --- | --- | --- | --- |
| 1 | Basmathi370 | 100 | 26 | VL Dhan16 | 100 | 51 | Sonasali | 60 |
| 2 | Dular** | 100 | 27 | AC41038 | 90 | 52 | Swarna | 60 |
| 3 | FR13A (LR) | 100 | 28 | MTU 1071 | 90 | 53 | WGL 347 | 60 |
| 4 | Jagannadh | 100 | 29 | NLR3242 | 90 | 54 | AC38460 | 50 |
| 5 | Konark | 100 | 30 | NLR34242 | 90 | 55 | Dikhow | 50 |
| 6 | Koshihikari | 100 | 31 | NLR4002 | 90 | 56 | ARC10533 | 40 |
| 7 | Minghui | 100 | 32 | Basmati<br>386 | 80 | 57 | Vasundhara | 40 |
| 8 | MTU1001 | 100 | 33 | BPT1235 | 80 | 58 | JGL3844 | 30 |
| 9 | MTU1010 | 100 | 34 | IR64 | 80 | 59 | NLR30491 | 30 |
| 10 | MTU1061 | 100 | 35 | JGL3855 | 80 | 60 | LN409 | 20 |
| 11 | MTU3626 | 100 | 36 | LN386 | 80 | 61 | Mahisugandh | 20 |
| 12 | N22** | 100 | 37 | NL42<sup>#</sup> | 80 | 62 | Shabigdhan | 20 |
| 13 | NBR16 | 100 | 38 | NLR3042 | 80 | 63 | Udayagiri | 20 |
| 14 | Nilagiri | 100 | 39 | Pokkali | 80 | 64 | BPT5204 | 10 |
| 15 | Nipponbare** | 100 | 40 | Rajeshwari | 80 | 65 | Dalasaitha | 10 |
| 16 | NL61<sup>#</sup> | 100 | 41 | Ranbir<br>Basmati | 80 | 66 | Disang | 10 |
| 17 | NL24<sup>#</sup> | 100 | 42 | WGL482 | 80 | 67 | Erramallelu | 10 |
| 18 | NLR145 | 100 | 43 | WGL915 | 80 | 68 | RNR150418 | 10 |
| 19 | NLR3238 | 100 | 44 | Kandagiri | 70 | 69 | Binuhungiri | 0 |
| 20 | Satya | 100 | 45 | Kolong | 70 | 70 | Kab Aus R270 | 0 |
| 21 | Siddi95 | 100 | 46 | IR1552 | 60 | 71 | Kapilee | 0 |
| 22 | Sona | 100 | 47 | NL1 | 60 | 72 | Lachit | 0 |
| 23 | Swarna Sub1A | 100 | 48 | NLR3354 | 60 | 73 | MTU1121 | 0 |
| 24 | TKM6 | 100 | 49 | NLR33671 | 60 | 74 | Pusa1121 | 0 |
| 25 | Vajram | 100 | 50 | NLR40024 | 60 |  |  |  |

\* - Survival percent (SP) under sub-lethal (SL) conditions, \*\* - reported heat tolerant genotypes, # - NL- NERICA Lines: derived from crosses involving *O. glaberrima*/*O. sativa* as parents. LR-Landrace.

A total of 11 genotypes were recorded in the class 0-20% SP (Figure 2) which means no germination at all or less germination, 6 genotypes were recorded in the class 20-40 % SP where germination is very less, 4 genotypes were recorded in the class 40-60% SP where the germination is a bit good, later 10 genotypes were recorded in the class of 60-80 % SP where the germination rate is good and a total of 43 genotypes were recorded in the class 80-100% SP where the germination rate is more and growth is observed in the genotypes.

**FIG 1.**
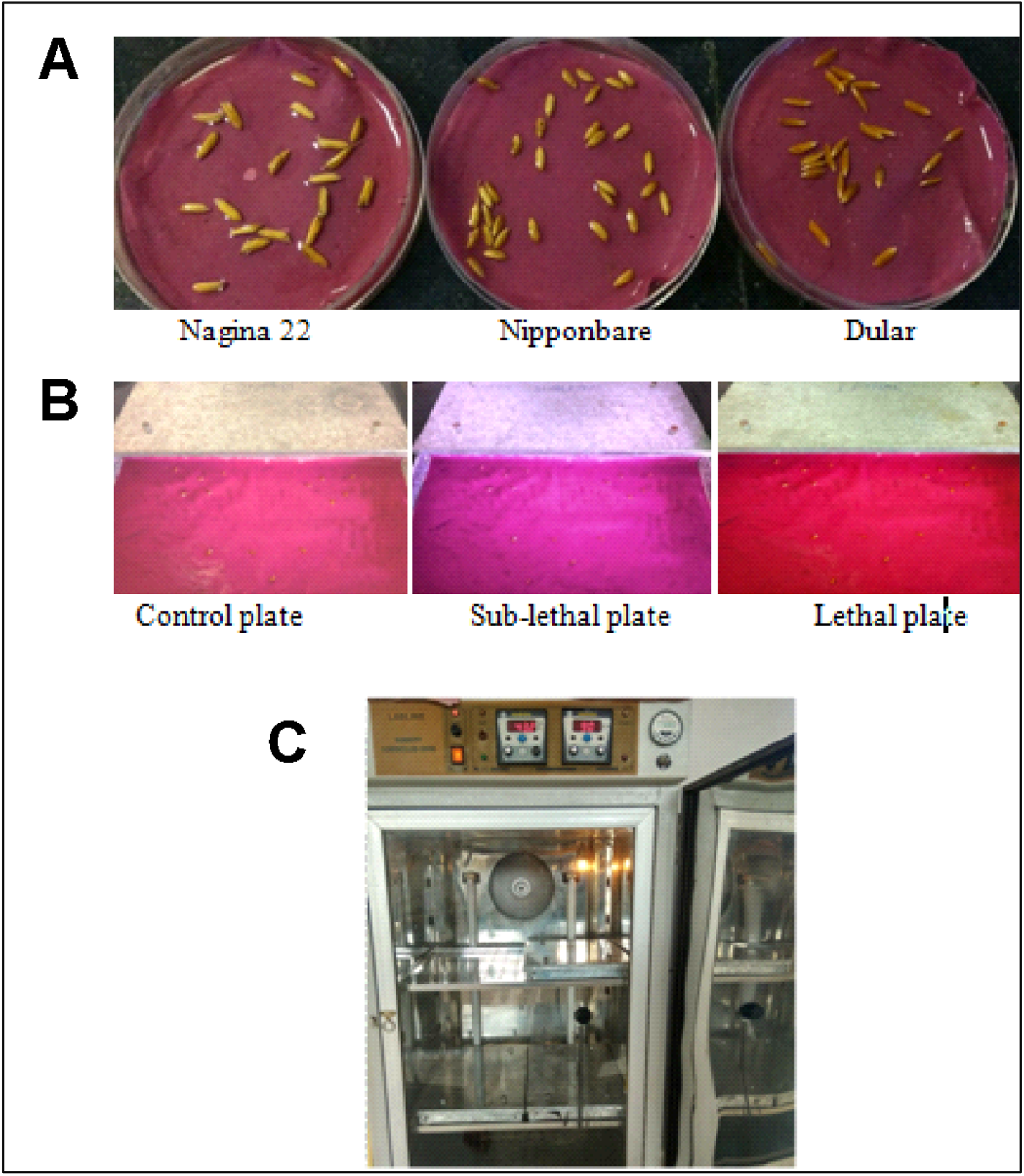
**A**. Germination of the three check genotypes N22, Dular and Nipponbare. **B**. The 3 check genotypes in Control, Sub-lethal (38-55°C) and lethal (55°C) plates. **C**. TIR chamber to induce high temperatures like Sub-lethal and lethal.

**FIG 2.**
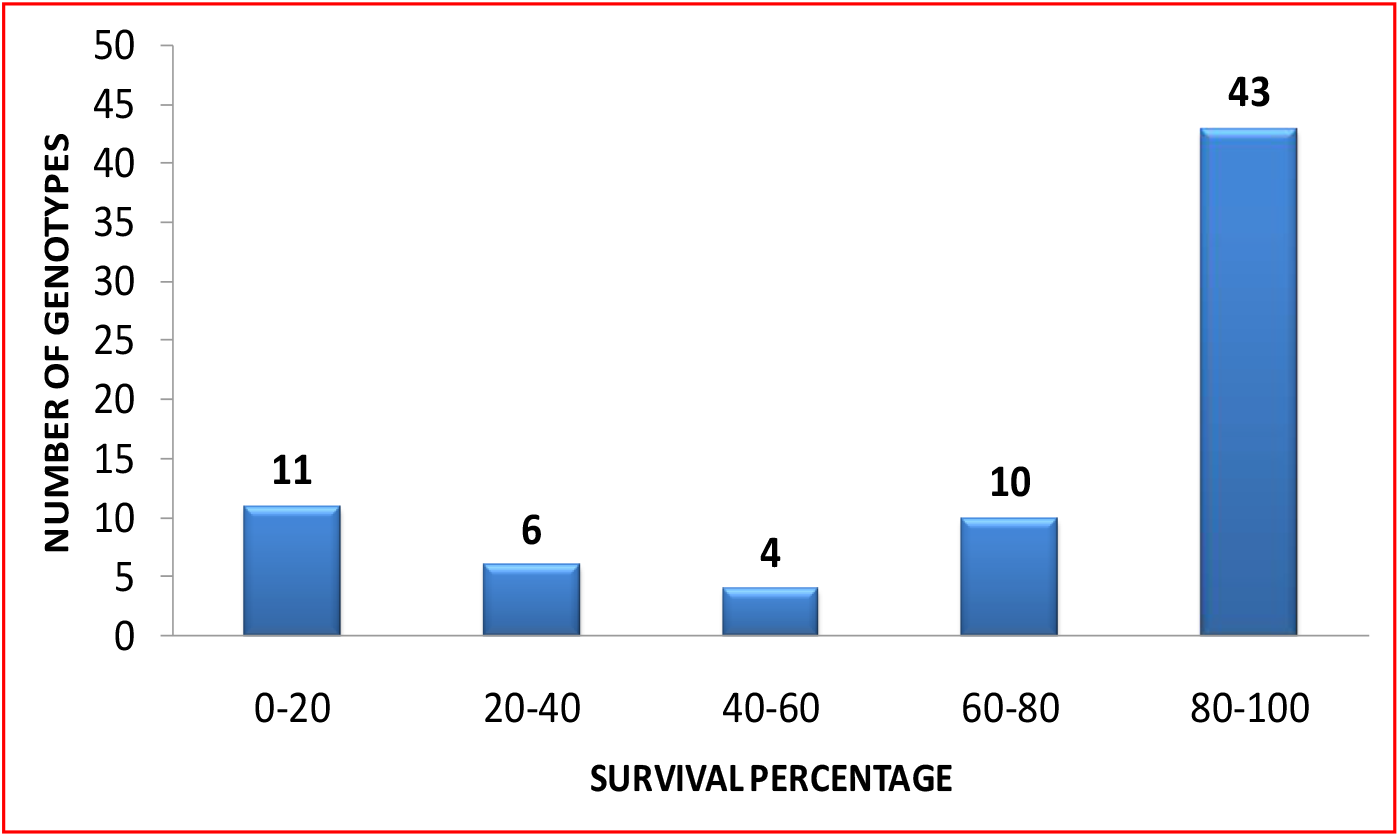
Phenotypic distribution pattern for survival percentage of genotypes under sub-lethal conditions

Certain phenotypic parameters were also taken to understand the survival percentage and growth of the survived seedlings post treatment under sub lethal conditions and compared with the control, which included maximum shoot length (cm) and root length (cm)s of both control and sub lethal genotypes (Table 3). Different parameters were taken into consideration to select the heat tolerant genotypes, Survival percentage, Maximum root lengths (MRL) of both control and sub lethal genotypes and also Maximum shoot lengths (SHTL) of both control and sub lethal genotypes (Table 3). The mean, standard deviation, skewness and kurtosis were calculated for all the genotypes under sub-lethal conditions and also for control plates. The skewness and kurtosis were maximum for MRL – control and SHTL-control with the values of 1.45, 1.72 and 8.54, 5.54 respectively (Table 3).

**TABLE 3.** Phenotypic parameters considered for heat tolerant genotypes.

| Trait | Mean | Minimum | Maximum | Standard Deviation | Skewness | Kurtosis |
| --- | --- | --- | --- | --- | --- | --- |
| Survival Percentage (%) | 67.57 | 0.00 | 100.00 | 35.14 | -1.58 | -1.64 |
| Maximum Root Length - Control (cm) | 6.92 | 2.64 | 14.24 | 1.47 | 1.45 | 8.54 |
| Maximum Root Length - Sub-Lethal (cm) | 5.71 | 0.70 | 10.74 | 2.76 | 0.01 | -0.73 |
| Shoot Length - Control (cm) | 7.17 | 3.46 | 14.42 | 1.78 | 1.72 | 5.54 |
| Shoot Length - Sub-Lethal (cm) | 5.30 | 1.72 | 11.75 | 2.03 | 0.85 | 0.61 |

A total of 52 genotypes were recorded in the class of 6.0-8.0 root length in cm in control and maximum of 22 genotypes were recorded in the class of 4.0-6.0 root length in cm in sub-lethal conditions (Table 4). Only one genotype was seen in maximum root length of 12.0-14.0 in control whereas 5 genotypes were observed in the root length class of 10.0-12.0 cm for sub-lethal conditions. Similarly, a maximum of 41 genotypes were considered under 6.0-8.0 shoot length in cm for control whereas 40 genotypes were recorded under 3.0-6.0 cm for shoot length for sub-lethal conditions (Table 4). Altogether only 2 genotypes were recorded in 14.0-16.0 cm for shoot length in control and a maximum of 5 genotypes were observed in 9.0-12.0 cm shoot length in sub lethal-conditions (Table 4).

**TABLE 4.** Performance of fourteen heat tolerant genotypes along with known check genotypes

| S No | Genotype | Survival Percentage (%) | Maximum Root Length-Control (cm) | Maximum Root Length - Sub-Lethal (cm) | Relative Root Length (cm) | Shoot Length-Control (cm) | Shoot Length-Sub-Lethal (cm) | Relative Shoot Length (cm) |
| --- | --- | --- | --- | --- | --- | --- | --- | --- |
| Check | N22 | 100 | 6.40 | 6.05 | -5.39 | 8.92 | 6.03 | -32.4 |
| Check | Nipponbare | 100 | 7.58 | 5.17 | -31.59 | 14.42 | 5.88 | -59.11 |
| Check | Dular | 100 | 7.40 | 10.61 | 43.36 | 7.58 | 6.33 | -16.57 |
| 1 | FR13A | 100 | 5.74 | 9.33 | 62.53 | 4.64 | 9.54 | 106.20 |
| **2** | Swarna Sub1A | 100 | 7.15 | 10.55 | 47.56 | 5.30 | 7.45 | 40.74 |
| **3** | NLR3238 | 100 | 7.23 | 10.22 | 41.36 | 6.33 | 8.44 | 33.38 |
| **4** | MTU1001 | 100 | 6.67 | 9.38 | 41.26 | 9.24 | 8.51 | -7.91 |
| **5** | Jagannadh | 100 | 7.48 | 10.58 | 41.74 | 10.39 | 9.33 | -10.16 |
| **6** | MTU1061 | 100 | 6.33 | 9.29 | 46.81 | 8.44 | 7.27 | -13.88 |
| **7** | Konark | 100 | 6.81 | 9.41 | 38.15 | 6.60 | 9.21 | 39.69 |
| **8** | Vajram | 100 | 5.27 | 8.48 | 61.00 | 9.24 | 5.23 | -43.40 |
| **9** | Satya | 100 | 5.21 | 5.80 | 11.31 | 6.31 | 6.16 | -2.33 |
| **10** | Minghui | 100 | 7.45 | 9.19 | 23.54 | 7.67 | 6.23 | -18.77 |
| **11** | VL Dhan16 | 100 | 4.45 | 4.80 | 7.87 | 5.26 | 7.44 | 41.39 |
| **12** | BPT1235 | 80 | 7.34 | 10.74 | 46.35 | 7.15 | 11.75 | 64.35 |
| **13** | JGL3855 | 80 | 6.50 | 9.26 | 42.67 | 6.64 | 7.25 | 9.21 |
| **14** | Basmathi386 | 80 | 7.11 | 8.51 | 19.77 | 6.60 | 9.49 | 43.82 |
|  | **Min** | **80** | **4.45** | **4.80** | **-31.59** | **4.64** | **5.23** | **-59.11** |
|  | **Max** | **100** | **7.58** | **10.74** | **62.53** | **14.42** | **11.75** | **106.20** |

### Heat tolerant genotypes

After checking for the survival percentage of the selected genotypes i.e., 43 which were recorded under 80-100% SP, only 14 were proven to show high root and shoot growth. Later it was found that FR13A and Swarna Sub 1 have performed well and gave good results than the already proven check Dular in terms of germination, survival, shoot length and root length (Table 4). All the 14 genotypes gave good performance at higher temperatures. Although 3 genotypes like BPT 1235, JGL 3855 and Basmati 386 showed 80% SP, the others showed 100% SP at sub-lethal temperatures (Table 4). It was observed that maximum root length is more for BPT 1235 (check) which is around 10.74 cm whereas it is less for VL Dhan 16 i.e., 4.80 in sub lethal conditions (Table 4). Also, it was observed that maximum shoot length is more for BPT 1235 i.e., around 11.75 cm and less for Vajram which is 5.23 in case of sub lethal conditions. In comparison of both the root and shoot lengths with control and sublethal the relativity is calculated i.e., relative lengths of root and shoot in which relative root length is maximum for FR 13A which is around 62.53 and it is minimum or very less for the check Nipponbare which is around −31.59 (Table 4). Also, the relative shoot length is maximum for FR 13A which is around 106.20 cm and very less for the check Nipponbare which is −59.11 proving that FR 13A is giving good results similar to that of the checks and compared with all the 14 varieties (Table 4).

The relative root and shoot lengths were calculated using maximum root and shoot lengths of control and sublethal. Out of all the three checks taken relative root and absolute shoot length is maximum in case of Dular whereas the relative root length and the relative shoot length of FR 13 A is maximum i.e., 62.53 and 106.20 respectively compared to other selected and tolerant varieties. Interestingly the values of FR13A were more than that of check variety Dular proving it to be highly tolerant.

### Molecular analysis

All the 51 primers were standardized between the temperatures 55°C-59°C (Table 5). Most of the primers were standardized at 59°C temperatures but very few were observed at 57°C which include RM10115, RM10469, RM15087, RM282, RM17270 and RM17296 (Table 5). It was also shown that the product size is maximum for RM19715 which is 350 bp and it was observed that the product size of the primer is less for RM16216 which is around 90-110 bp (Table 5). We designed a set of 8 primers with respect to heat tolerance in rice using Primer 3 software (Table 6). These primers were designed from different HSP families and TT 1 (Thermal tolerance gene) gene targeted markers. We chose four polymorphic primers namely, RM16216, RM 17271, RM 5687 and a TT 1 gene targeted marker, TTC/ TTM primers in the study (Table 5 and Table 6) for screening all the 14 genotypes each from heat tolerant and sensitive groups to understand the allelic patterns.

**TABLE 5.** Detailed list of reported SSR primers, their annealing temperature and their observed product sizes (bp)

| S.No. | Primer Name | Chr | Annealing temperature | Product Size (bp) |
| --- | --- | --- | --- | --- |
| 1 | RM10021 | 1 | 59 | 120 |
| 2 | RM10115 | 1 | 57 | 100 |
| 3 | RM10469 | 1 | 57 | 200 |
| 4 | RM157 | 1 | 59 | 150 |
| 5 | RM323 | 1 | 59 | 250 |
| 6 | RM490 | 1 | 59 | 100 |
| 7 | RM183 | 2 | 59 | 300 |
| 8 | RM5631 | 2 | 59 | 100 |
| 9 | RM324/RM424/RM13021 | 2 | 59 | 100 |
| 10 | RM15077 | 3 | 59 | 200 |
| 11 | RM15080 | 3 | 59 | 280 |
| 12 | RM15081 | 3 | 59 | 100 |
| 13 | RM15087 | 3 | 57 | 190 |
| 14 | RM16088 | 3 | 59 | 200 |
| 15 | RM16184 | 3 | 59 | 200 |
| 16 | RM16199 | 3 | 59 | 250 |
| 17 | RM16216 | 3 | 59 | 90-110 |
| 18 | RM282 | 3 | 57 | 150 |
| 19 | RM554 | 3 | 59 | 250 |
| 20 | RM16490 | 4 | 59 | 300 |
| 21 | RM16530 | 4 | 59 | 100 |
| 22 | RM16838 | 4 | 59 | 200 |
| 23 | RM16933 | 4 | 59 | 100 |
| 24 | RM17270 | 4 | 57 | 190-210 |
| 25 | RM17296 | 4 | 57 | 350 |
| 26 | RM17496 | 4 | 59 | 280 |
| 27 | RM17544 | 4 | 59 | 200 |
| 28 | RM5687 | 4 | 59 | 80-120 |
| 29 | RM18034 | 5 | 59 | 150 |
| 30 | RM274 | 5 | 59 | 180 |
| 31 | RM19341 | 6 | 59 | 300 |
| 32 | RM19429 | 6 | 59 | 300 |
| 33 | RM19715 | 6 | 59 | 350 |
| 34 | RM16029 | 8 | 59 | 200 |
| 35 | RM547 | 8 | 59 | 100 |
| 36 | RM 24303 | 9 | 59 | 100 |
| 37 | RM24571 | 9 | 59 | 300 |
| 38 | RM24596 | 9 | 59 | 100 |
| 39 | RM24631 | 9 | 59 | 150 |
| 40 | RM566/RM24303 | 9 | 59 | 100 |
| 41 | RM25649 | 10 | 59 | 150 |
| 42 | RM25654 | 10 | 59 | 150 |
| 43 | RM26212 | 11 | 59 | 250 |

**TABLE 6.** List of genic primers, primer annealing temperature and their observed product sizes (bp)

| S.No. | Primer Name | Locus | Chr | Tm | Product Size (bp) |
| --- | --- | --- | --- | --- | --- |
| 1 | RGNMS264 | LOC_Os01g43650 | 1 | 59 | 100 |
| 2 | TTC/TTW | LOC_Os03g26970 | 3 | 59 | 130 |
| 3 | TTC/TTM | LOC_Os03g26970 | 3 | 59 | 100 and 350 |
| 4 | RGNMS2015 | LOC_Os05g45410 | 5 | 59 | 200 |
| 5 | RGNMS2168 | LOC_Os06g11610 | 6 | 59 | 250 |
| 6 | SVHT801 | LOC_Os02g42801 | 10 | 59 | 115 |
| 7 | RGNMS3524 | LOC_Os11g32030 | 11 | 59 | 350 |
| 8 | SVHT665 | LOC_Os12g07665 | 12 | 59 | 200 |

Several primers were selected from the literature based on their performance with respect to heat tolerance in several rice varieties. The product size of the primers were between 90 - 350 where the maximum size is observed in case of RM 19715 and minimum price is observed for RM 16216.

The allelic patterns of the polymorphic markers were observed between checks, tolerant and sensitive genotypes (Fig 3). Several genotypes exhibited allelic patterns between 80-120 bp in case of tolerant group but not 120bp (Fig 3) with the primer RM5687.

**FIG 3.**
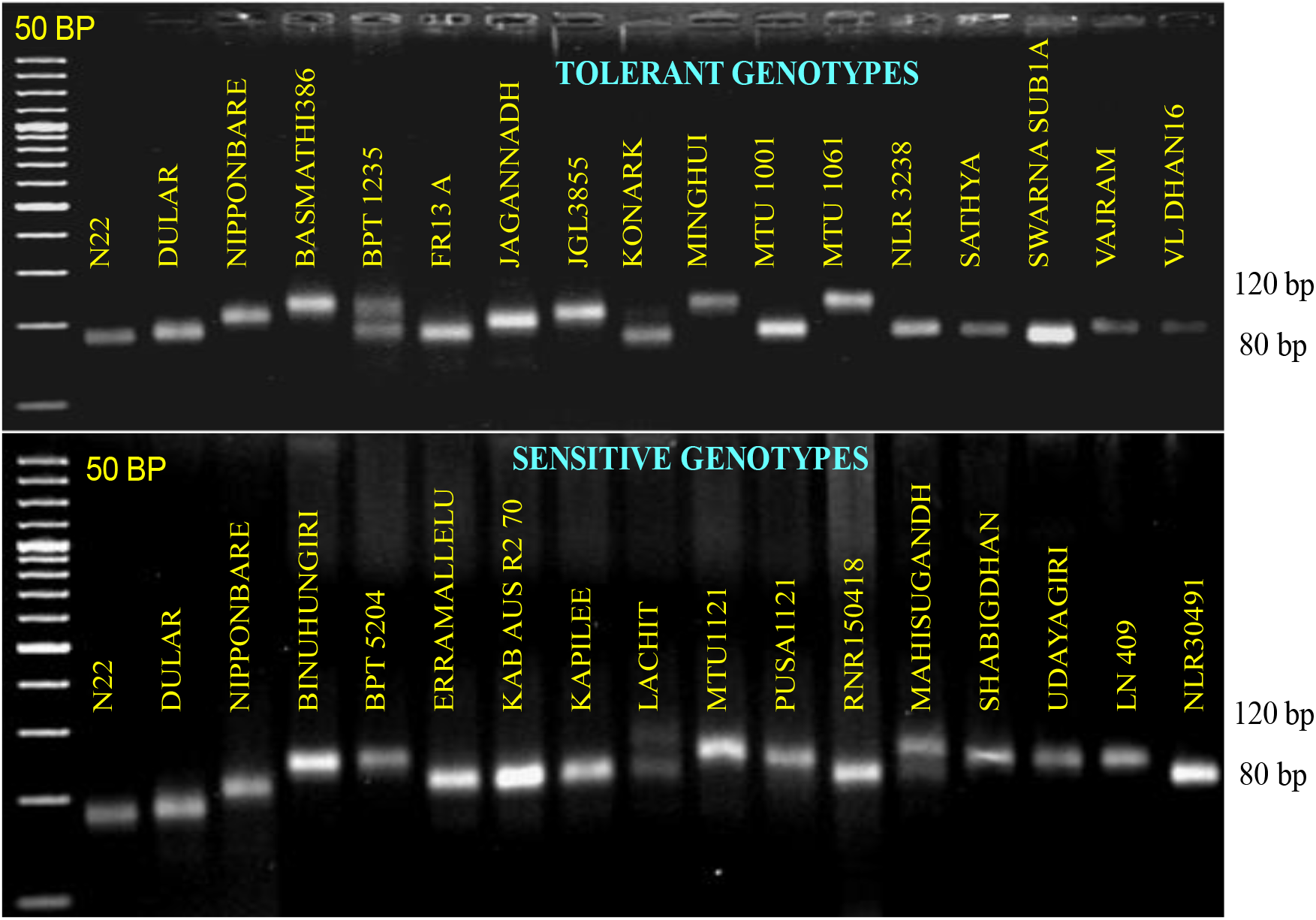
Polymorphic amplification pattern generated by RM5687 marker between the selected heat tolerant and sensitive genotypic classes under study

Further, it was determined that both RM 17270 and RM 5687 were heat tolerant primers with polymorphic patterns, and also the distribution of genotypes between checks using these primers was understood (Table 7). The three major varieties like Basmati 386, FR13A, Swarna Sub1A were concluded to have high heat tolerance not only due to their phenotypic characters but also their polymorphic patterns were also considered with the checks. Basmati 386 gave less growth compared to the other two varieties when considered along with the phenotypic characters (Table 7).

**TABLE 7.** Genotypes that showing similar allele pattern with reported heat tolerant sources for markers RM17270 and RM5687

| Genotypes showing similar allele pattern as reported heat tolerant check |  |  |  |  |
| --- | --- | --- | --- | --- |
| PRIMERS | N 22* | DULAR* | NIPPONBARE* | DULAR*/<br>NIPPONBARE* |
| RM 17270 | - | - | - | Basmati 386, FR13A,<br>Swarna Sub1A |
| RM 5687 | Konark | Sathya, Vajram,<br>VLDhan16,<br>Erramallelu#, Kab<br>AusR270# | Jaganadh,<br>Binuhungiri#, BPT<br>5204#, Lachit#,<br>Udayagiri#, LN 409# | - |

\*- reported heat tolerant genotypes, #- heat sensitive genotypes

The TTC/TTM primer of TT 1 gene showed the polymorphic patterns as the other primers obtained from the literature. Mostly all the genotypes gave good results showing heat tolerance because alleles are seen near to above 350 bp and few are less than that i.e., near 100 bp alleles (Fig.4). The alleles that were obtained by different polymorphic markers were not clearly in distinguishing the tolerant set from sensitive set of genotypes. Among the reported genotypes also the allele sizes were varied except for the marker TTC/TTM, as they were from different sub groups of rice (Fig. 4). N22 and Dular comes under *Aus* group, whereas Nipponbare comes under *japonica* group. To understand this clearly allele codes were assigned for each primer.

**FIG 4:**
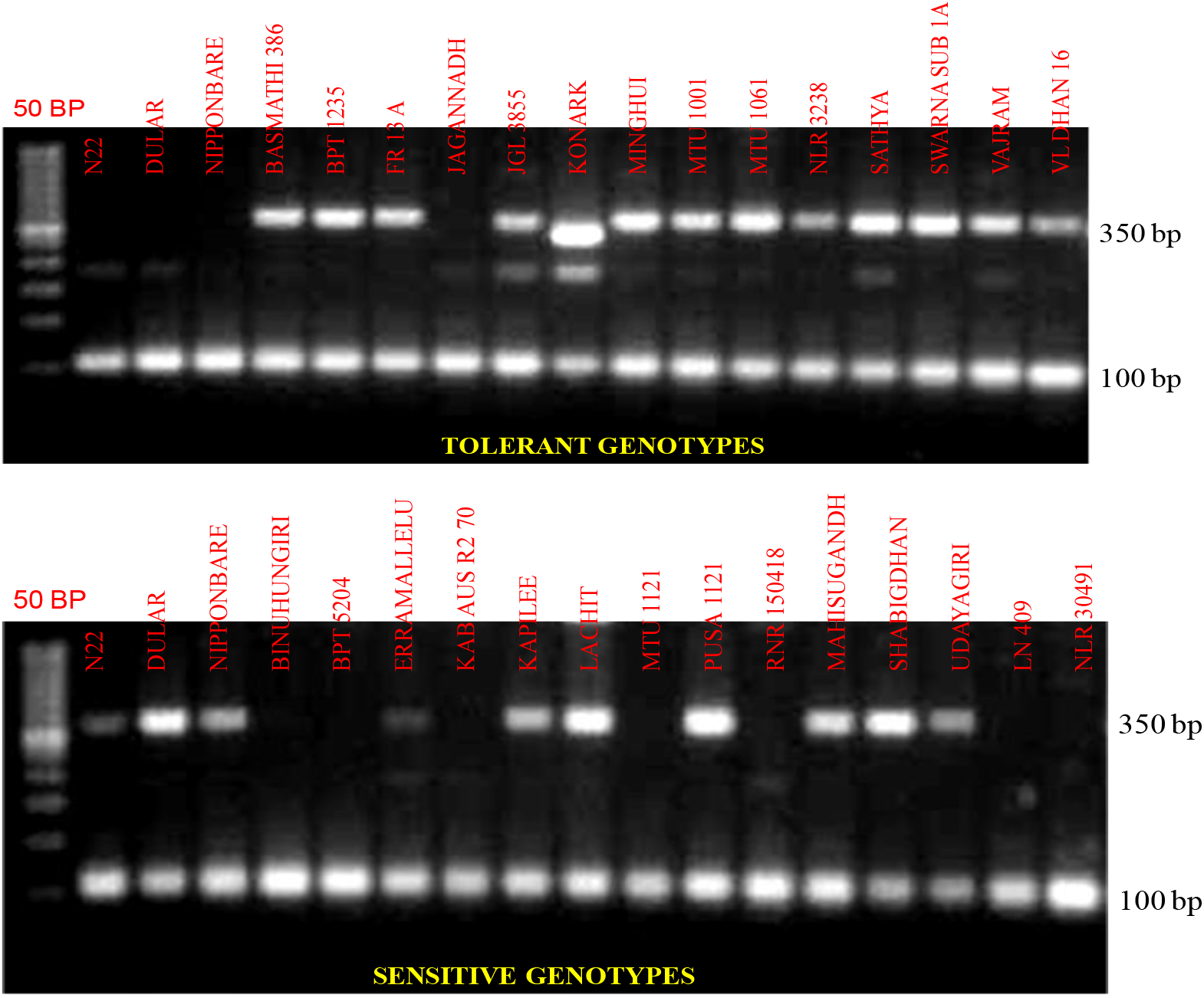
Polymorphic amplification pattern of TTC/TTM primer with tolerant and sensitive genotypes used in the present study.

**FIG 5:**
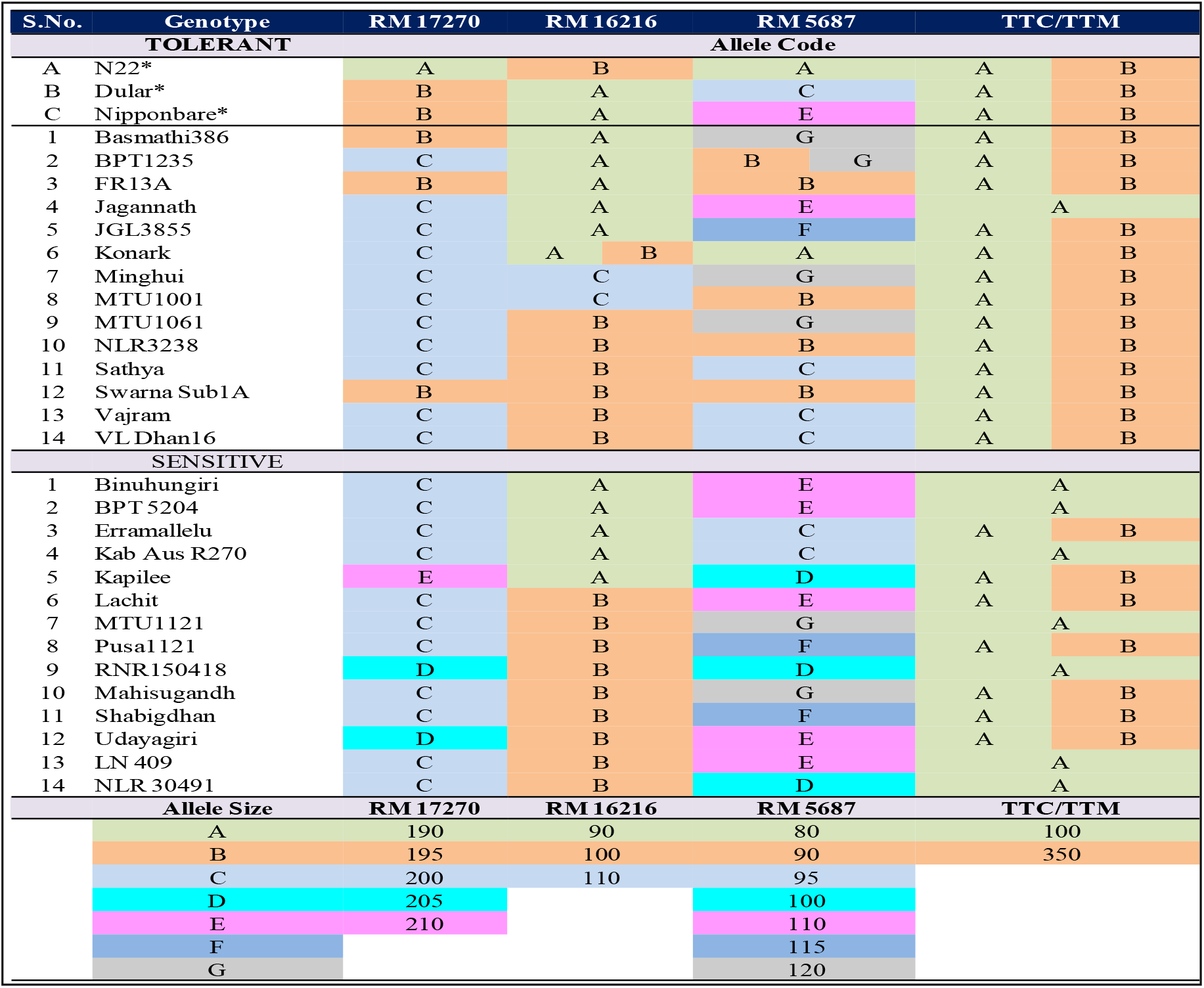
Allele score as a check to detect heat tolerant varieties.

### Analysing the informativeness of polymorphic markers

Each genotype was assigned an allele code generated by respective polymorphic marker based on their allele size and colour coded as shown in Figure 6. As many as a highest number of 100bp alleles were identified by TTC/TTM in the range of 100 to 350bp (Table 7). It was understood that the marker RM17270 was amplified at 190bp in N22, but no other genotype showed similar allele size to N22. The same marker was amplified at 195bp in Dular and Nipponbare genotypes, wherein Basmathi386, FR13A and Swarna Sub1A were also shown similar amplification pattern like Dular and Nipponbare (Table 7). The marker RM16216 showed three kinds of alleles i.e. 90bp, 100bp and 110bp. Of which, 90bp and 100bp are major alleles (Table 7). By considering the allele patterns and allele codes it was proved that all the 14 genotypes from the tolerant set showed maximum polymorphic alleles (Table 7). Hence, the identified list of tolerant genotypes from the study, especially FR13A, Swarna Sub1A, Sathya, Vajram, VLDhan16, Basmathi386 and Jagannadh can be used for further gene identification studies apart from targeted breeding programmes (Table 7).

## DISCUSSIONS

High temperature stress (increase in more than tolerable air temperature) is perhaps the main natural element impacting crop development, advancement, and yield measures. This pressure initiates numerous biochemical, molecular and physiological changes and reactions that impacts different cells and entire plant measures that influence crop yield and quality (Wei *et al*., 2012). The impact of environmental stresses, particularly those of dry season and heat stress, has been studied separately. However, under natural conditions, both of these stresses often occur in union (Jagadish *et al*., 2010). The worldwide expansion in temperature will likewise expand the seriousness of other natural pressing factors like floods and dry seasons. Varieties in precipitation will cause flooding and more incessant dry seasons which are the main deterrents for the internal waters and vigorous planting (Veerasekaran *et al*., 2021).

The present investigation was, therefore, undertaken to study the effect of high temperature on survivability, growth and its component traits on different rice genotypes, to identify heat tolerant genotypes and to validate and tag the reported molecular markers and as well newly designed genic SSR markers among high temperature progressive and vulnerable genotypes. Also, genetic diversity is the foundation of the genetic improvement of crop plants as it serves as a reservoir for identifying superior lines that can withstand heat stress (Liang *et al*., 2010; Wanwarang *et al*., 2020). Selective Line Genotyping (SLG) used in this study involves initial polymorphism study with extreme few numbers (two) of genotypes to allow/score remaining alleles (allelic variants) of high-types with polymorphic markers identified using extreme core bulks. Later we used those identified polymorphic markers to screen 14 genotypes each that fall in both extreme phenotypic classes individually (Yongyoa *et al*., 2019). Thus, SLG can be believed to be superior to BLA, as it arrives at identification of more kinds of alleles and allele patterns of genotypes; and also, their degree of trait association at particular locus. This is evident through different degrees of occurrence of each allele among high-types in the current study (Katherine *et al*., 2020). Therefore, polymorphic primers are considered very important to understand tolerant and sensitive genotypes because of their high informativeness (Ye *et al*., 2012).

Consequential variations exist among rice germplasm in response to maximum temperature stress *viz*. blooming at cooler occasions of day (early daytime blossoming), more dust feasibility, bigger anthers, longer basal dehiscence and presence of long basal pores, are some of the phenotypic markers for high temperature tolerance (Kheir *et al*., 2012). So, the development of more sustainable, resilient agricultural systems could be achieved by identifying heat tolerant rice genomes and development of new rice varieties for mitigating the yield losses under high temperature conditions during *summer* and *kharif* seasons (Kobayashi *et al*., 2011). The temperature above 35°C results in low seed setting rate, causes floret sterility and abnormal pollination (Lin *et al*., 2012). Day by day temperatures, higher than 30°C or every day greatest temperatures higher than 35°C during the blossoming time frame will bring about helpless anther dehiscence.

Plants’ heat response is highly complex (Hedhly *et al*., 2011). In current days, some reputed techniques like gene editing are being used to address the heat tolerant/induced functional basis of the genotype. From the recent experiments conducted, it was proven that high temperature decreases the grain filling period in basmati rice from 32 to 26 days, reduced yield by 6%, and caused a decrease in absolute starch (3.1%) and amylose content (22%). Quantifiable exercises of key chemicals associated with sucrose to starch change, sucrose synthase, ADP-glucose pyrophosphorylase, starch phosphorylase and dissolvable starch synthase in endosperms created at 32°C were lower than those at 22°C contrasted and comparable aging stage on an endosperm premise. Specifically, granule-bound starch synthase (GBSS) action was fundamentally lower than comparing action in endosperms creating at 22°C during every formative stage (Ahmed *et al*., 2015).

The genotypes selected for the study include genotypes from Indica, Japonica Javanicas, Aus groups and wild relatives of rice. These genotypes are selected and are allowed to grow under sub–lethal conditions. The genotypes giving good performance i.e., survivability, maximum root and shoot lengths are selected and are separated from the entire genotype sets (Alberio *et al*., 2018). High temperatures are induced by using TIR protocol in which temperatures to a maximum level and control in humidity can be done (Sudhakar *et al*., 2013). TIR technique relies on genotype’s acquired tolerance, to determine their heat tolerance, wherein a gradual exposure of genotypes to a temperature regime over a period was used. Using TIR technique, it was proved that sufficient genetic variability was present among rice genotypes for high temperature tolerance. The percent survival of seedlings varied from 0 to 100 % with a mean of 80.33%. The percent reduction in root growth varied from 0 to 73 % with a mean of 20.89 %. Four genotypes namely NLR 34242, NLR40066, NLR40059, NLR40050 also exhibited higher thermotolerance without reduction in root but slight reduction in seedling survival by 10%. Similar studies were also conducted by Vijaylakshmi *et al*. (2015)

Later molecular analysis was carried out to check the polymorphic patterns obtained by heat tolerance specific SSR primers. A total of 51 primers were selected out of which 43 are chosen from literature and 8 are designed by using Primer 3 software, these designed primers belong to different HSP (Heat Shock protein) families (Guo *et al*., 2020). Molecular markers are very important tools in the understanding the genetic variation and in interpretation of genetic relationships within and among different species (Zhu *et al*., 2017). Simple sequence repeats (SSRs), also known as the microsatellite DNA marker, is an effective tool for identifying genetic differences of germplasm resources with the advantages of ease, clarity and maximum polymorphism, co-dominance and its steadiness (Liang *et al*., 2010, Ma *et al*., 2010). There has been developing interest with the use of marker-based models in recent years. In the studies conducted by using these models, descriptions of the effect of allele coding system on inference and computations are often absent or missing (Lv *et al*., 2017). Also, several other common allele coding alters these regression coefficients by deducting a value from each marker such that the mean of regression coefficients is zero within each marker. It is also known as centered allele coding (Stranden *et al*., 2011). This kind of coding is highly useful to identify the genotypes that fall within similar allele size ranges for a major number of loci, to identify like allele pattern with proven genotypes (heat tolerant) and to group them under a class (Shim *et al*., 2020).

Thus, keeping in lieu of the above identified potently reported polymorphic set of markers, irrespective of their allelic differences among the genotypes it can be concluded that the efficiency of selective line genotyping technique used in the current study is remarkable in associating effective markers to the trait (Sharma et al., 2019; 2020). Further the efficiency of the technique is well explored with limited genotyping panel to arrive at small number of useful markers out of huge tests, for further studies (Gouda *et al*., 2020). The above outcomes propose that the TIR strategy is an amazing and productive method to recognize hereditary changeability in high temperature resistance in rice inside a brief timeframe and it is appropriate for screening an enormous number of genotypes (Hsuan *et al*., 2019).

## CONCLUSION

Rice occupies 23% of the total area under cereal production in the world. It is the staple nourishment for the greater part of the total populace with Asia addressing the biggest producer and consumer region. For the most part, rice is unfavourably influenced by high temperature in the lower heights of the tropical regions. Under molecular analysis carried out in the present study, a set of 51 reported and as well new SSR and genic markers were employed to assess the genetic diversity among the 74 rice genotypes. These markers gave good amplification with prominent alleles at 57°C and 59°C. By employing Selective line genotyping approach; both of each fourteen selected Tolerant and Sensitive genotypes along with three reported genotypes (N22, Dular and Nipponbare) were subjected for genotyping with a total of 51 markers. Out of 51 markers 92% were monomorphic between heat tolerant and sensitive classes, as well as compared to checks. Four markers i.e., RM17270, RM16216, RM5687, TTC/TTM, were polymorphic. Despite the fact that, allele coding is a viable strategy the alleles produced from various polymorphic markers were not unmistakably recognized lenient set from touchy arrangement of genotypes like TIR procedure. The recognized 14 genotypes of rice can be utilized as contributor hotspot for growing high temperature lenient rice genotypes to oppose worldwide ascent temperature.

## Notes

### Competing Interest Statement

The authors have declared no competing interest.

## REFERENCES

1. Ahmed, N., Tetlow, I. J., Nawaz, S., Iqbal, A., Mubin, M and Rehman, M. S., 2015. Effect of high temperature on grain filling period, yield, amylose content and activity of starch biosynthesis enzymes in endosperm of basmati rice. Journal of Science, Food and Agriculture 95, 2237–2243.

2. Bado S, Forster BP, Mukhtar AGA, Cieslak JJ, Berthold G, Luxiang L (2016) Protocols for pre-field screening of mutants for salt tolerance in rice, wheat and barley. Springer open access, ISBN 978-3-319-26590-2

3. Burke, J.J., (2011). Identification of genetic diversity and mutations in higher plant acquired thermotolerance. Physiol.Plant. 112:167–170.

4. C. Alberio, L.A. Aguirrezábal, N.G. Izquierdo, R. Reid, S. Zuil, A. Zambelli Effect of genetic background on the stability of sunflower fatty acid composition in different high oleic mutations J Food Agric Environ, 98 (11) (2018), pp. 4074–4084

5. Chaturvedi, A.K., Bahuguna, R.N., Shah, D. et al. High temperature stress during flowering and grain filling offsets beneficial impact of elevated CO2 on assimilate partitioning and sink-strength in rice. Sci Rep 7, 8227 (2017). https://doi.org/10.1038/s41598-017-07464-6

6. FAO. (2014). FAO statistical year book. FAO of the United Nations, Rome, pp 1171, 76.

7. FAO. (2016). Rice Market Monitor. Volume XIX Issue No. 1. Pp 1–44 www.fao.org/economic/RMM.

8. Fu Y., Gu Q., Dong Q., Zhang Z., Lin C., Hu W., Pan R., Guan Y., Hu J. Spermidine enhances heat tolerance of rice seeds by modulating endogenous starch and polyamine metabolism. Molecules. 2019;24:1395. doi: 10.3390/molecules24071395.

9. Gouda, A.C., Ndjiondjop, M.N., Djedatin, G.L. et al. Comparisons of sampling methods for assessing intra- and inter-accession genetic diversity in three rice species using genotyping by sequencing. Sci Rep 10, 13995 (2020). https://doi.org/10.1038/s41598-020-70842-0.

10. Guo, LM., Li, J., He, J. et al. A class I cytosolic HSP20 of rice enhances heat and salt tolerance in different organisms. Sci Rep 10, 1383 (2020). https://doi.org/10.1038/s41598-020-58395-8

11. Hedhly, A.,et al.,2011. Sensitivity of flowering plant gametophytes to temperature fluctuations. Environment Experimental Botany 74:9–16.

12. Hsuan, TP., Jhuang, PR., Wu, WC. et al. Thermotolerance evaluation of Taiwan Japonica type rice cultivars at the seedling stage. Bot Stud 60, 29 (2019). https://doi.org/10.1186/s40529-019-0277-7

13. Huang X.M., Zhao Y., Wei X., Li C., Wang A., Zhao Q., Li W., Guo Y., Deng L., Zhu C., et al. Genome-wide association study of flowering time and grain yield traits in a worldwide collection of rice germplasm. Nat. Genet. 2012;44:32–39. doi: 10.1038/ng.1018.

14. Jagadish S, Muthurajan R, Oane R, Wheeler T, Heuer S, Bennett J, Craufurd Q (2010). Physiological and proteomic approaches to address heat tolerance during anthesis in rice. J. Exp. Bot. 61: 143–156.

15. Jagadish, S. V. K., Septiningsih, E. M., Kohli, A., Thomson, M. J., Ye, C., Redoña, E., et al. (2012). Genetic advances in adapting rice to a rapidly changing climate. J. Agron. Crop Sci. 198, 360–373. doi: 10.1111/j.1439-037X.2012.00525.x

16. Jain, B., Sarial, A., Kaushik, P., 2019. Understanding G× E interaction of elite basmati rice (Oryza sativa L.) genotypes under north indian conditions using stability models. Applied Ecology and Environmental Research 17, 5863–5885.

17. Jain, B., Sarial, A., Kaushik, P., others, 2018. Stability analysis utilizing AMMI model and regression analysis for grain yield of Basmati rice (Oryza sativa L.) genotypes. Journal of Experimental Biology and Agricultural Sciences 6, 522–530.

18. Katherine., S, Tulloch, M.Q., Burns, M. et al. Developing KASP Markers for Identification of Basmati Rice Varieties. Food Anal. Methods 14, 663–673 (2021). https://doi.org/10.1007/s12161-020-01892-3.

19. Kesh, H., Kaushik, P., 2020. Impact of Marker Assisted Breeding for Bacterial Blight Resistance in Rice: A Review. Plant Pathology Journal 19, 151–16.

20. Kesh, H., Kharb, R., Ram, K., Munjal, R., Kaushik, P., Kumar, D., others, 2021. Adaptability and AMMI biplot analysis for yield and agronomical traits in scented rice genotypes under diverse production environments. Indian Journal of Traditional Knowledge (IJTK) 20, 550–562.

21. Kheir MS, Sheshshayee T, Prasad G, Udayakumar M (2012) Temperature induction response as a screening technique for selecting high temperature-tolerant cotton lines. J Cotton Sci 16:190–199

22. Kim J., Shon J., Lee C.K., Yang W., Yoon Y., Yang W.H., Kim Y.G., Lee B.W. Relationship between grain filling duration and leaf senescence of temperate rice under high temperature. Field Crops Res. 2011;122:207–213. doi: 10.1016/j.fcr.2011.03.014.

23. Kobayashi KM, Mayuko YY (2011). Percentage of dehisced thecae and length of dehiscense control pollination stability of rice cultivars at high temperatures. Plant Production Science. 14: 89–95.

24. Krishnan P., Ramakrishnan B., Reddy K.R., Reddy V.R. Advances in Agronomy. Volume 111. Academic Press; London, UK: 2011. High-temperature effects on rice growth, yield, and grain quality; pp. 87–206.

25. Kumar, A., Kaushik, P., 2021. Heat Stress and its Impact on Plant Function: An Update. Preprints.

26. Kumar, S.P.J., Susmita, C., Agarwal, D.K. et al. Assessment of Genetic Purity in Rice Using Polymorphic SSR Markers and Its Economic Analysis with Grow-Out-Test. Food Anal. Methods 14, 856–864 (2021). https://doi.org/10.1007/s12161-020-01927-9.

27. Lafarge T., Bueno C., Frouin J., Jacquin L., Courtois B., Ahmadi N. Genome-wide association analysis for heat tolerance at flowering detected a large set of genes involved in adaptation to thermal and other stresses. PLoS ONE. 2017;12:e0171254. doi: 10.1371/journal.pone.0171254.

28. Liang, J., Yan, L., Peng, X., Mei, S., Harold, C and Bao, J. S., 2010. Genetic diversity and population structure of a diverse set of rice germplasm for association mapping. Theor Applied Genetics 121:475–87.

29. Lin HY, Wu YP, Hour AL, Ho SW, Wei FJ, Hsing YI and Lin YR (2012). Genetic diversity of rice germplasm used in Taiwan breeding programs. Bot. Studies. 53: 363–376

30. Liu QH, Wu X, Li T, Ma JQ, Zhou XB (2013) Effects of elevated air temperature on physiological characteristics of flag leaves and grain yield in rice. Chilean J Agric Res 73(2):85–90

31. Liu S, Waqas MA, Wang S, Xiong X, Wan Y (2017) Effects of increased levels of atmospheric CO2 and high temperatures on rice growth and quality. PLoS ONE 12(11):e0187724, pp 1–15

32. Lv, Q., Huang, Z., Xu, X. et al. Allelic variation of the rice blast resistance gene *Pid3* in cultivated rice worldwide. Sci Rep 7, 10362 (2017). https://doi.org/10.1038/s41598-017-10617-2

33. Ma, Q., Dai, X., Xu, Y., Guo, J., Liu, Y., Chen, N., Xiao, J., Zhang, D., Xu, Z., Zhang, X., and Chong, K., 2010. Enhanced tolerance to chilling stress in OsMYB3R-2 transgenic rice is mediated by alteration in cell cycle and ectopic expression of stress genes. Plant Physiology. 150, 244–256.

34. Malhi, G.S., Kaur, M., Kaushik, P., 2021. Impact of climate change on agriculture and its mitigation strategies: A review. Sustainability 13, 1318.

35. Prasanth VV, Chakravarthi DVN, Vishnu KT, Venkateswara RY, Panigrahy M, Mangrauthia SK (2012) Evaluation of rice germplasm and introgression lines for heat tolerance. Ann Biol Res 3:5060–5068

36. Priyanka, V., Goel, N., Dhaliwal, I., Sharma, M., Kumar, R., Kaushik, P., 2021a. Epigenetics: A Key to Comprehending Biotic and Abiotic Stress Tolerance in Family Poaceae.

37. Priyanka, V., Kumar, R., Dhaliwal, I., Kaushik, P., 2021b. Germplasm Conservation: Instrumental in Agricultural Biodiversity-A Review.

38. Raza A, Razzaq A, Mehmood SS, et al. Impact of Climate Change on Crops Adaptation and Strategies to Tackle Its Outcome: A Review. Plants (Basel). 2019;8(2):34. Published 2019 Jan 30. doi:10.3390/plants8020034

39. S. Zhu, R.L. Huang, H.P. Wai, H.L. Xiong, X.H. Shen, H.H. He, S. Yan, Mapping quantitative trait loci for heat tolerance at the booting stage using chromosomal segment substitution lines in rice. Physiol Mol Biol Plants, 23 (2017), pp. 817–825

40. Sajeevani Weerasekara, Clevo Wilson, Boon Lee & Viet-Ngu Hoang (2021) Impact of natural disasters on the efficiency of agricultural production: an exemplar from rice farming in Sri Lanka, Climate and Development, DOI: 10.1080/17565529.2021.1893635

41. Sathishraj R., Bheemanahalli R., Ramachandran M., Dingkuhn M., Muthurajan R., Jagadish S.V.K. Capturing heat stress induced variability in spikelet sterility using panicle, leaf and air temperature under field conditions. Field Crops Res. 2016;190:10–17. doi: 10.1016/j.fcr.2015.10.012.

42. Sato H., Todaka D., Kudo M., Mizoi J., Kidokoro S., Zhao Y., Shinozaki K., Yamaguchi-Shinozaki K. The Arabidopsis transcriptional regulator DPB 3-1 enhances heat stress tolerance without growth retardation in rice. Plant Biotech. J. 2016;14:1756–1767. doi: 10.1111/pbi.12535.

43. Shah F, Huang J, Kul K, Nie L, Shah T, Chen C (2011) Impact of high-temperature stress on rice plant and its traits related to tolerance. J Agric Sci 149:545–556. https://doi.org/10.1017/S0021859611000360

44. Shanmugavadivel, P. S., Mithra, A. S., Prakash, C., Ramkumar, M. K., Tiwari, R., Mohapatra, T., et al. (2017). High resolution mapping of QTLs for heat tolerance in rice using a 5K SNP array. Rice 10:28. doi: 10.1186/s12284-017-0167-0

45. Sharma, V., Saini, D.K., Kumar, A., Kaushik, P., 2019. A Review of Important QTLs for Biofortification Traits in Rice.

46. Sharma, V., Saini, D.K., Kumar, A., Kesh, H., Kaushik, P., 2020. Breeding for Biofortification Traits in Rice: Means to Eradicate Hidden Hunger. Agronomy-Climate Change & Food Security 35.

47. Shi W., Xiao G., Struik P.C., Jagadish K.S., Yin X. Quantifying source-sink relationships of rice under high night-time temperature combined with two nitrogen levels. Field Crops Res. 2017;202:36–46. doi: 10.1016/j.fcr.2016.05.013.

48. Shim, KC., Kim, S.H., Lee, HS. et al. Characterization of a New *qLTG3–1* Allele for Low-temperature Germinability in Rice from the Wild Species *Oryza rufipogon*. Rice 13, 10 (2020).https://doi.org/10.1186/s12284-020-0370-2.

49. Stranden,I., Christensen,O.F.,2011. Allele coding in genomic evaluation. Genetics Selection Evolution:43–25.

50. Sudhakar,P.,Venkatesh,B.d.,Sarath, K.R.Y.,2013. Screening of thermotolerant ragi genotypes at seedling stage using tir technique. Journal of Life Sciences the Bioscan 8(4): 1493–1495.

51. Tenorio FA, Ye C, Redona E, Sierra S, Laza M, Argayoso liang(2013) Screening rice genetic resources for heat tolerance. SABRAO J Breed Genet 45:371–381

52. Vallejos, C. E. (2007). An Expedient and Versatile Protocol for Extracting High-Quality DNA From Plant Leaves. New York, NY: Cold Spring Harbor Protocols.

53. Vijaylakshmi,D., Srividhya,S.,Vivitha,P., Ravindran,M.,2015. Temperature induction response (TIR) as a rapid screening protocol to dissect the genetic variability in acquired thermotolerance in rice and to identify novel donors for high temperature stress tolerance. Indian Journal of Plant Physiology.Vol 20:368–374.

54. Wanwarang.P., Natjaree.P., Kannika.S. (2020). Genetic diversity and allelic frequency of selected Thai and exotic rice germplasm using SSR markers. Science direct. 6 393–403.

55. Wei-hun Z, Da-wie X, Guo-ping Z (2012) Identification and physiological characterization of thermo-tolerant rice genotypes. J Zhejiang Univ, Agri & Life Sci 38(1):1–9

56. Xie, L., Tan, Z., Zhou, Y., Xu, R., Feng, L., Xing, Y., et al. (2014). Identification and fine mapping of quantitative trait loci for seed vigor in germination and seedling establishment in rice. J. Int. Plant Biol. 56, 749–759. doi: 10.1111/jipb.12190

57. Ye, C., Argayoso, M. A., Redoña, E. D., Sierra, S. N., Laza, M. A., Dilla, C. J., et al. (2012). Mapping QTL for heat tolerance at flowering stage in rice using SNP markers. Plant Breed. 131, 33–41. doi: 10.1111/j.1439-0523.2011.01924.x

58. Yongyoa Xie.,Jintao.T.,Jianle.H.2019. An asymmetric allelic interaction drives allele transmission bias in rice hubrids. Nature communications. 10..2501 (2019).

59. Yu K., Chen G., Patrick W.H., Jr. Reduction of global warming potential contribution from a rice field by irrigation, organic matter, and fertilizer management. Glob. Biogeochem. Cycles. 2014;18 doi: 10.1029/2004GB002251.

60. Yufang Xu., Chengcai Chu., Shankguo Yao., The impact of high temperature stress on rice: Problems and solutions. The Crop Journal.https://doi.org/10.1016/j.cj.2021.02.011.

